# Cryo-EM structures of an anti-MLC1 Fab in apo and peptide-bound states reveal the structural basis of antigen recognition

**DOI:** 10.1101/2025.10.08.681078

**Authors:** Junmo Hwang, Kunwoong Park, Jeong-Sun Choi, Tae-Ryong Riew, Kyung-Ok Cho, Kyoung Tai No, Hyun-Ho Lim

**Affiliations:** Neurovascular Unit Research Group, Korea Brain Research Institute (KBRI), Daegu 41062, Republic of Korea; Structural Biology Division, Baobab AiBIO Co., LTD., Incheon, 21984, Republic of Korea; Department of Pharmacology, College of Medicine, The Catholic University of Korea, Seoul 06591, Republic of Korea; Department of Anatomy, College of Medicine, The Catholic University of Korea, Seoul 06591, Republic of Korea; Department of Medical Science, College of Medicine, Graduate School of The Catholic University of Korea, Seoul 06591, Republic of Korea; Institute of Convergence Science and Technology, Yonsei University, Incheon 21983, Republic of Korea; Department of Brain Sciences, DGIST, Daegu 42988, Republic of Korea

**Keywords:** MLC1, Monoclonal antibody 37E5, Fab fragment, Cryo-electron microscopy, Epitope-paratope interaction, Induced-fit mechanism

## Abstract

Monoclonal antibodies are indispensable tools in structural biology and biomedical research, but defining the molecular basis of their specificity remains challenging. Here, we developed a novel monoclonal antibody (37E5) against the astrocytic membrane protein MLC1, a component of gliovascular signaling implicated in megalencephalic leukoencephalopathy with subcortical cysts. 37E5 demonstrated high specificity and versatility across biochemical, cellular, and histological assays, enabling reliable detection of MLC1 in both human and mouse tissue. Using single-particle cryo-EM, we determined ∼3 Å resolution structures of the 37E5 Fab in apo and antigen-bound states, despite the small molecular mass (∼50 kDa), close to the lower size limit of cryo-EM. The antigen-bound structure revealed continuous density for an MLC1-derived peptide and enabled atomic mapping of polar and non-polar interaction networks. Conformational changes in CDR-L1 and CDR-L2 indicated an induced-fit mechanism of recognition. Comparison with AlphaFold-predicted models underscored the accuracy of Fab backbone prediction but revealed major limitations in modeling epitope-paratope geometry. These findings establish 37E5 as a versatile antibody for mechanistic studies of gliovascular biology and MLC disease, while demonstrating that cryo-EM can achieve atomic-level characterization of small Fab-antigen complexes, thereby expanding the methodological frontier of antibody-antigen structural biology.

**Significance:** This study defines the molecular basis of MLC1 recognition by a novel monoclonal antibody, establishes 37E5 as a versatile reagent for mechanistic and translational research, and demonstrates the feasibility of cryo-EM to resolve dynamic features of small Fab-antigen complexes.

## MAIN

Monoclonal antibodies are indispensable tools in therapeutics, diagnostics, and modern biological research owing to their high specificity and affinity for target antigens. By recognizing and modulating antigen function, they provide versatile means to dissect molecular mechanisms and develop effective interventions. Antigen recognition is mediated by complementarity-determining regions (CDRs) within the variable domains of heavy and light chains, and understanding the structural basis of these epitope–paratope interactions is critical for antibody engineering and functional applications.

X-ray crystallography has long been the gold standard for determining antibody–antigen complex structures. More recently, cryo-electron microscopy (cryo-EM) has emerged as a powerful alternative, particularly for complexes that are flexible or refractory to crystallization. Although high-resolution cryo-EM has traditionally been limited by the molecular mass of macromolecules, recent studies have reported near-atomic structures of proteins below 100 kDa, including alcohol dehydrogenase (82 kDa), methemoglobin (64 kDa), streptavidin (52 kDa), a riboswitch (40 kDa), CDK-activating kinase (85 kDa), human serum albumin (65 kDa), maltose binding protein (43 kDa), and the kinase domain of human PLK1 (37 kDa)^1-7^. Nevertheless, high-resolution cryo-EM analysis of whole IgG molecules (∼150 kDa) remains challenging due to hinge flexibility^8^, and even Fab fragments—although widely used as surrogates—are relatively small (∼50 kDa) and contain an intrinsically flexible elbow region that can hinder particle alignment and limit attainable resolution^9^.

Recent progress has demonstrated that these barriers can be overcome. For example, the epitope region (∼20 kDa) of human carcinoembryonic antigen-related cell adhesion molecules in complex with tusamitamab (tusa Fab) was resolved at ∼3.1 Å, revealing atomic details of the epitope– paratope interface^10^. Furthermore, by introducing disulfide bonds at the elbow region, engineered rigid-Fab complexes with small antigens (21 and 26 kDa) were determined at high resolutions of 2.3∼2.5 Å by single-particle cryo-EM^11^. Despite these advances, structural determination of apo Fab fragments has so far been limited to resolutions below ∼6 Å^12^, and peptide-bound Fab structures have not yet been reported.

MLC1 is an astrocytic membrane protein that forms complexes with GlialCAM at perivascular endfeet, where it contributes to ion and water homeostasis in the brain and peripheral blood cells^13-15^. Mutations in MLC1 or GlialCAM cause megalencephalic leukoencephalopathy with subcortical cysts (MLC), a childhood-onset leukodystrophy characterized by macrocephaly, seizures, and white matter vacuolation. Despite its clinical importance, the molecular architecture and functional role of MLC1 remain poorly understood. This knowledge gap has been further compounded by the absence of validated antibodies that are cross-reactive across species and broadly applicable in diverse experimental settings, limiting both mechanistic studies and assay development.

To address these challenges, we developed a novel monoclonal antibody against MLC1, 37E5, and rigorously evaluated its immunoreactivity across multiple contexts, including different MLC1 constructs, orthologs, heterologously expressed protein in HEK293 cells, endogenous protein from mouse brain lysates, and fixed mouse and human brain samples. These analyses, performed using western blotting, co-immunoprecipitation, and immunofluorescence staining, demonstrated robust and specific recognition of MLC1. In addition, protein–protein interactions between the 37E5 antibody or its Fab fragment and purified human MLC1 were assessed by size-exclusion chromatography, further confirming their direct association. Together, these results establish 37E5 as a highly specific antibody suitable for diverse experimental applications in the detection and characterization of MLC1.

To gain molecular insight into the specific interaction between the 37E5 antibody and MLC1, we determined 3.16 Å and 3.04 Å cryo-EM structures of the Fab in both apo form and in complex with an MLC1-derived peptide, respectively. The antigen-bound structure revealed salt bridges and hydrogen bonding networks between the epitope and complementarity-determining regions (CDRs). Comparison with the apo Fab uncovered an induced-fit rearrangement of the CDRs upon antigen engagement. These findings define the structural basis of MLC1 recognition and provide a framework for engineering improved antibodies and developing epitope-tag systems for protein purification and detection.

## RESULTS

### Development and validation of a novel monoclonal antibody against MLC1, 37E5

Despite the clinical and biological importance of MLC1, progress in mechanistic studies has been hampered by the lack of validated antibodies suitable for biochemical and structural analyses. To address this limitation, we generated a monoclonal antibody using an N-terminal 14–amino acid peptide derived from mouse MLC1. Hybridoma screening identified 37E5 as the lead clone with high binding activity^16^ (Extended Data Fig. 1). We next assessed its specificity across multiple experimental contexts. In heterologous expression systems, 37E5 recognized both human and mouse MLC1 (hMLC1 and mMLC1) with N-terminal Flag tags expressed in HEK293 cells with strong and reproducible signals in western blotting. The specificity was confirmed by western blotting with anti-Flag tag antibody. Both 37E5 and anti-Flag antibodies clearly showed monomeric, dimeric, and trimeric forms of MLC1 as previously reported^17,18^ (Fig. 1a). 37E5 is also able to precipitate both hMLC1 and mMLC1 proteins in the detergent-solubilized cell lysate (Fig. 1b). We further validated the ability of 37E5 to detect endogenous MLC1 by performing immunoprecipitation followed by western blotting with mouse brain lysates. This analysis demonstrated that the antibody recognizes native protein in a complex tissue background (Fig. 1c). These results indicate that 37E5 can efficiently pull down MLC1 and is suitable for probing protein-protein interactions involving MLC1.

**Figure 1.**
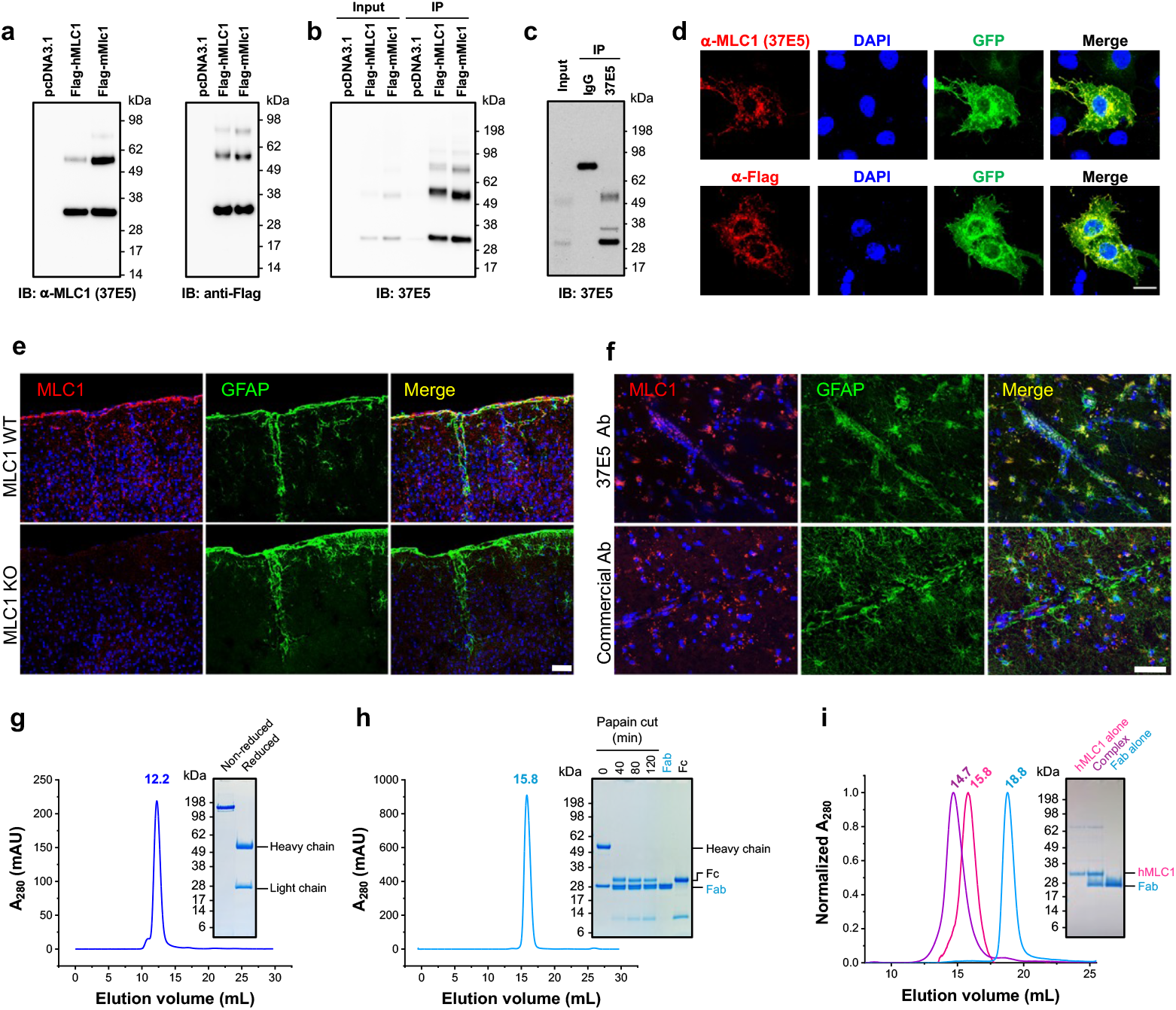
Development and validation of the anti-MLC1 monoclonal antibody 37E5. (a) Schematic illustration of the strategy used to generate the monoclonal antibody 37E5 against human MLC1. (b) Western blot analysis of HEK293 cells expressing MLC1 constructs and orthologs. 37E5 specifically detected human MLC1, with no cross-reactivity observed in non-transfected controls. (c) Immunoprecipitation of endogenous MLC1 from mouse brain lysates using 37E5, followed by western blot detection. The antibody efficiently precipitated native MLC1 from complex tissue extracts. (d) Immunofluorescence staining of Flag::hMLC1::GFP-transfected HEK293 cells with 37E5. Signals were observed at the plasma membrane and in intracellular compartments, overlapping with GFP fluorescence and consistent with the localization of overexpressed MLC1. Control staining with anti-Flag antibody produced comparable results. Scale bar, XX μm. (e) Immunofluorescence staining of cortical sections from wild-type mouse brain showing robust labeling of astrocytic processes surrounding blood vessels. In contrast, no staining was detected in Mlc1 knockout mouse brain, confirming the antibody’s specificity. Scale bar, XX μm. (f) Immunofluorescence staining of human hippocampal brain sections with 37E5. The antibody revealed distinct astrocytic membrane staining similar to the distribution observed in mouse brain. Scale bar, XX μm. (g) SDS-PAGE analysis of purified 37E5 IgG obtained from hybridoma cultures. (h) SDS-PAGE analysis of Fab fragments generated by papain digestion of 37E5. (i) Size-exclusion chromatography showing direct binding between recombinant MLC1 and 37E5 Fab fragments, providing biochemical confirmation of the antibody–antigen interaction.

Furthermore, 37E5 demonstrated robust performance in immunofluorescence staining of fixed cells and brain tissues. In *Flag::hMLC1::GFP*-transfected HEK293 cells (Fig. 1d), 37E5 produced signals at the plasma membrane as well as within intracellular organelles, which closely overlapped with GFP fluorescence, consistent with the expected localization of overexpressed MLC1. Control experiments with an anti-Flag antibody produced staining patterns comparable to those obtained with 37E5. Having established its specificity in transfected cells, we next investigated whether 37E5 could detect endogenous MLC1 in brain tissue. In wild-type mouse brain cortical sections (Fig. 1e), 37E5 prominently labeled astrocytic processes surrounding blood vessels, consistent with the known enrichment of MLC1 at perivascular endfeet^14,19-22^. In contrast, no staining was observed in Mlc1 knockout mouse brain^23^, confirming the antibody’s specificity. The perivascular staining pattern in wild-type tissue was consistent with the expected gliovascular distribution and demonstrated that 37E5 effectively recognizes endogenous MLC1 in complex tissue environments. Similarly, in human hippocampal brain sections (Fig. 1f), 37E5 produced distinct astrocytic membrane staining that closely resembled the distribution observed in mouse brain. The ability of 37E5 to detect MLC1 across species and in both cellular and histological contexts underscores its specificity and versatility, establishing it as a reliable reagent for biochemical, cellular, and tissue-based assays.

To evaluate the antigen-binding capacity of the 37E5 antibody and Fab fragment, the antibody was purified from hybridoma cultures and digested with papain to generate Fab fragments (Fig. 1g, h). Size-exclusion chromatography demonstrated direct binding between recombinant MLC1 carrying different affinity tags and the full-length 37E5 antibody (Extended Data Fig. 2), as well as between recombinant MLC1 and the Fab fragments (Fig. 1i). These results provide additional biochemical evidence supporting the specificity of the antibody–antigen interaction. Together, these results establish 37E5 as a highly specific monoclonal antibody with broad applicability in biochemical, cellular, and histological assays. The availability of this reagent addresses a long-standing limitation in MLC1 research and provides a foundation for structural studies and mechanistic dissection of MLC disease biology.

### High-resolution cryo-EM structures of apo and antigen-bound 37E5 Fab

To elucidate the molecular determinants of antigen-antibody specificity, we determined single-particle cryo-EM structures of the 37E5 Fab in both apo and antigen-bound states. Cryo-EM grids prepared with purified Fab or Fab incubated with an MLC1-derived peptide yielded high-quality micrographs. Data collection and image processing resulted in reconstructions at 3.16 Å and 3.04 Å resolution for the apo and antigen-bound states, respectively, despite the relatively small size of the Fab (∼50 kDa), which approaches the lower size limit for high-resolution cryo-EM analysis (Extended Data Table 1; Extended Data Fig. 3 and 4). These results underscore the feasibility of resolving near-atomic structures of small antibody fragments by cryo-EM (Fig. 2a, b). Following 3D volume reconstruction, atomic models were built using ModelAngelo^24^. The amino acid sequences of the Fab variable regions were inferred from DNA sequences obtained by multiplex RT-PCR, as described previously^25^. The constant region sequences were derived from ModelAngelo-generated models and refined by manual inspection (Extended Data Fig.5).

**Figure 2.**
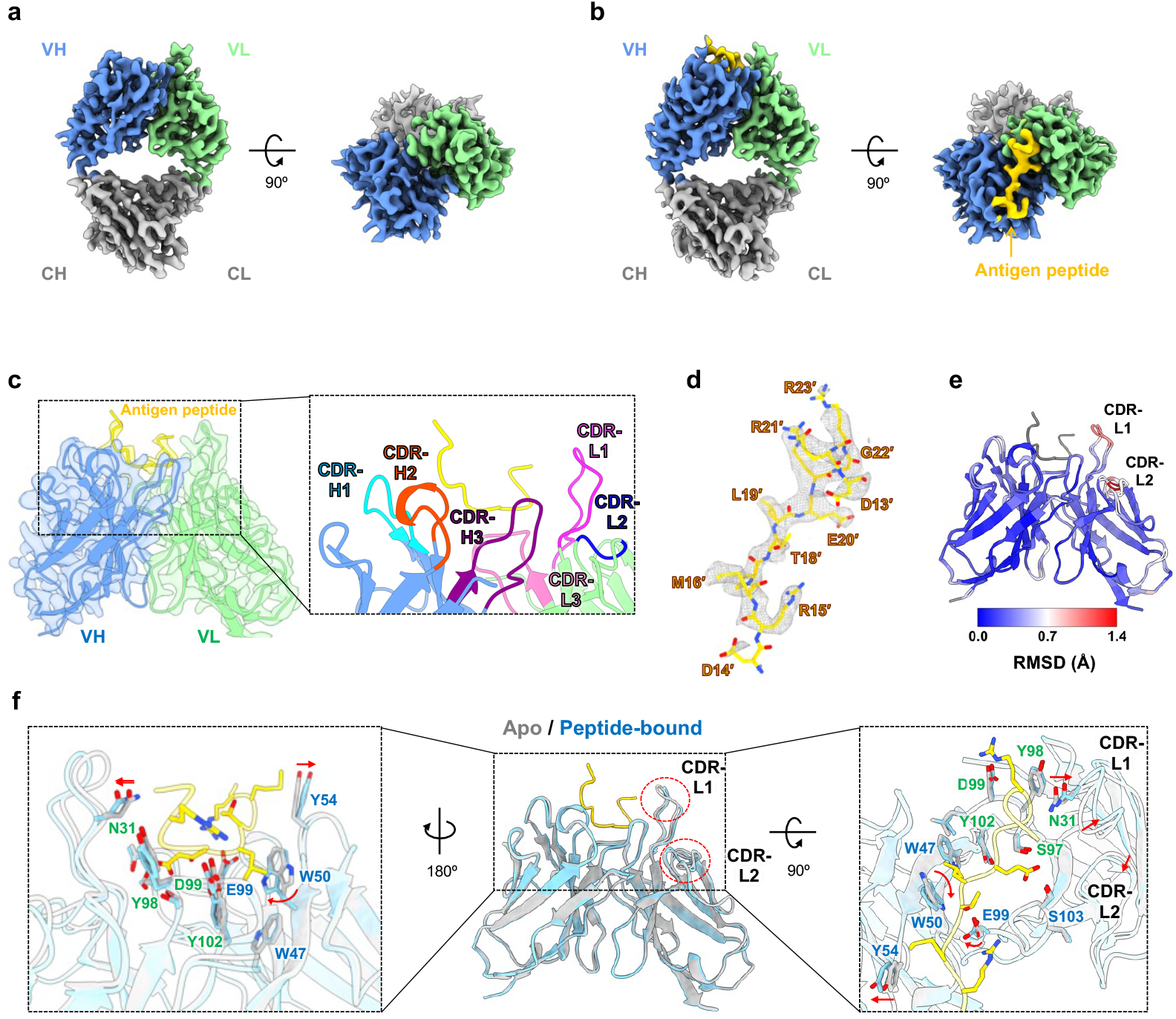
Cryo-EM structures of apo and antigen-bound 37E5 Fab. (a) Cryo-EM density map of the apo 37E5 Fab reconstructed at ∼3.X Å resolution (a) and the antigen peptide–bound Fab complex (b). The overall VH and VL domain architecture is clearly resolved. (c) Structural model of the Fab–peptide complex fitted into the cryo-EM map, highlighting the antigen-binding cleft formed by the CDR loops. (d) Close-up atomic model of the MLC1-antigen peptide fitted into the cryo-EM map. (e) Cα RMSD highlights local rearrangements in the CDR loops upon antigen engagement. (f) Structural changes between apo and peptide-bound Fab illustrating antigen-induced conformational motions of the CDR loops. Two rotational views are shown to emphasize the induced-fit rearrangements that stabilize the antigen-binding interface.

Both the apo and antigen-bound Fab reconstructions revealed the canonical organization of the Fab fragment, consisting of two constant (CL and CH) and two variable (VH and VL) domains, and showed well-defined density for the framework regions and the complementarity-determining regions (CDRs) (Fig. 2a, b). In the antigen-bound complex, additional continuous density corresponding to the antigen peptide (*yellow*, Fig. 2b) was observed at the paratope, enabling unambiguous placement of the peptide within the binding cleft. Detailed inspection of the binding pocket revealed a paratope formed by all six CDR loops (CDR-L1, CDR-L2, CDR-L3, CDR-H1, CDR-H2, and CDR-H3) (Fig. 2c). The high local resolution in this region allowed unambiguous side-chain assignment for both antibody and antigen, enabling precise mapping of protein–protein interactions (Extended Data Fig. 6 and 7). The peptide density itself was continuous and well resolved (Fig. 2d), permitting accurate modeling of key residues critical for binding specificity.

Comparison of the apo and peptide-bound structures revealed pronounced conformational changes at the antigen-binding site. RMSD analysis of Cα atoms highlighted local rearrangements in the CDRs upon peptide binding, particularly in CDR-L1 and CDR-L2, which underwent clear conformational shifts (Fig. 2e). Conformational transition analysis further illustrated these changes, demonstrating how the CDR loops reorient and stabilize in the bound state, consistent with an induced-fit mode of antigen recognition (Fig. 2f). Upon peptide binding, Trp50 and Glu99 underwent side-chain rotations, while Asn31 and Tyr54 shifted outward to create space for the antigen. Concomitantly, the outward displacement of CDR-L1 contributed to stabilizing the binding interface (Fig. 2f).

Together, the cryo-EM structures define the overall architecture of the 37E5 Fab and provide direct visualization of the MLC1 peptide epitope within the paratope. They also reveal antigen-induced conformational changes that stabilize the binding interface. These findings establish the molecular basis of MLC1 recognition and demonstrate the power of cryo-EM to resolve dynamic structural features even in small (∼50 kDa) antibody–antigen complexes.

### Structural determinants of epitope–paratope specificity in MLC1 recognition

To define the molecular determinants of 37E5–MLC1 recognition, we first examined the interaction networks within the binding pocket. Polar contacts formed an extensive network of salt bridges and hydrogen bonds between the antigen peptide and the Fab CDRs (Fig. 3a). Residue Arg15′ of MLC1 formed a salt bridge with Glu99 in the VH domain, while Arg23′ interacted with Asp99 in the VL domain. Residues Glu20′ and Gly22′ established hydrogen-bonding networks with Ser103 in VH as well as Tyr102, Ser97, Tyr98, and Asn31 in VL. Throughout this work, peptide residue numbering follows the full-length mouse MLC1 sequence and is denoted with an apostrophe. These polar interactions were complemented by non-polar contacts: Met16′ and Leu19′ of MLC1 engaged Tyr54, Trp47, and Trp50 in VH and Leu100 in VL (Fig. 3b). The hydrophobic side chains of the peptide were accommodated by a groove within the paratope, thereby stabilizing the orientation of the bound peptide.

**Figure 3.**
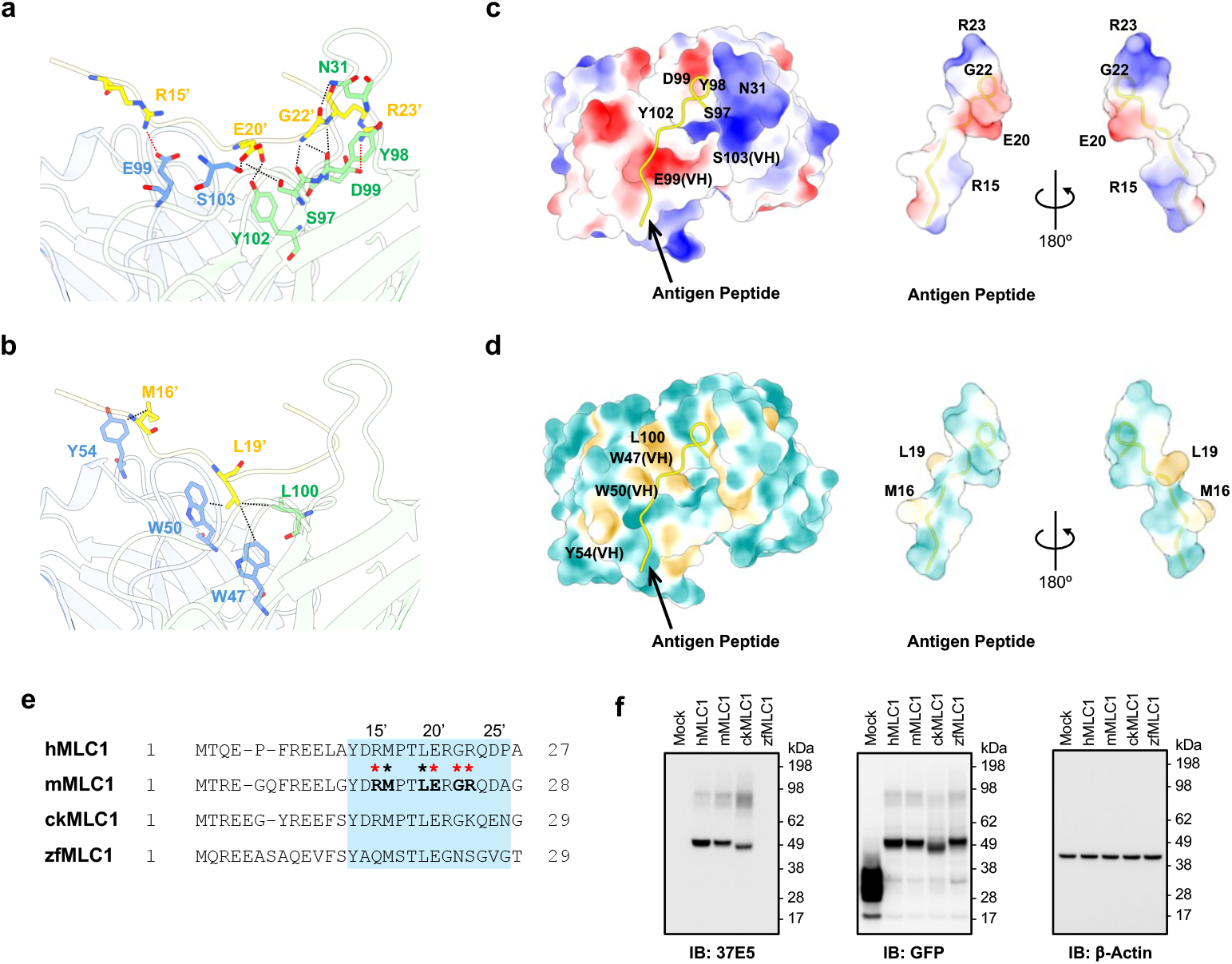
Structural and functional determinants of 37E5–MLC1 specificity. (a) Polar interaction network between the MLC1 peptide and the Fab paratope, highlighting hydrogen bonds and salt bridges involving key residues (e.g., Arg15, Gly22, Arg23). (b) Non-polar interaction network showing hydrophobic contacts that stabilize the peptide within the binding cleft. (c, d) Surface property maps of the Fab–peptide interface. Electrostatic surface potentials reveal clear charge complementarity between the Fab paratope and the antigen peptide (c), whereas hydrophobicity mapping highlights non-polar clusters that contribute to peptide anchoring (d). The antigen peptide is shown as a ribbon model on the Fab surface (*left*), and its surface properties are illustrated separately (*right*). (e) Sequence alignment of the MLC1 epitope region among human, mouse, chicken, and zebrafish orthologs. Asterisks indicate residues involved in epitope–paratope interactions (*red*, polar; *black*, non-polar). (f) Western blot validation of MLC1 orthologs fused to C-terminal GFP and expressed in HEK293 cells. 37E5 detected strong signals in human and mouse, a reduced signal in chicken, and no signal in zebrafish. Anti–β-actin served as a loading control, and anti-GFP confirmed expression of all constructs.

Surface property analysis further revealed a high degree of complementarity between the two binding partners. Electrostatic mapping showed that the positively charged surface of the MLC1 peptide aligned with negatively charged regions of the CDRs, while hydrophobicity analysis highlighted a cluster of non-polar residues anchoring the peptide within the binding pocket (Fig. 3c and 3d). These surface properties provide a structural explanation for the high specificity of 37E5 binding.

The structural determinants of specificity were further supported by evolutionary and functional analyses. Sequence alignment of MLC1 orthologs showed that residues critical for binding (Arg15′, Met16′, Leu19′, Glu20′, Gly22′, and Arg23′) were conserved in human and mouse, whereas Arg23′ was substituted by Lys in chicken, and Arg15′, Gly22′, and Arg23′ were replaced by Gln, Asn, and Ser, respectively, in zebrafish (Fig. 3e; Extended Data Fig. 1). Western blotting of GFP-tagged MLC1 orthologs expressed in HEK293 cells confirmed these predictions: 37E5 strongly recognized human and mouse MLC1, produced moderately reduced signals for chicken, and showed no detectable binding to zebrafish MLC1 (Fig. 3f). These results show that epitope residue substitutions explain the species-dependent recognition pattern of 37E5, with conservation in mammals supporting strong binding, and divergent residues in chicken and zebrafish reducing or abolishing reactivity.

Taken together, these structural, biophysical, and biochemical analyses identify the polar and non-polar interaction networks, electrostatic complementarity, and conserved epitope residues as key determinants of 37E5–MLC1 specificity. These findings highlight how structural insight, integrated with biochemical and evolutionary data, can explain antibody specificity and provide a framework for rational antibody design, enabling the precise modulation of specificity and binding affinity.

### Experimental validation reveals limitations of AI-predicted antibody–antigen models

To assess the utility of computational prediction for antibody–antigen complexes, we compared our cryo-EM structure of the 37E5 Fab–peptide complex with an AlphaFold-predicted model. Overall alignment revealed that the Fab backbone was predicted with high accuracy, showing close agreement in the VH, VL, CH, and CL domains (Fig. 4a). This result is consistent with previous reports that AlphaFold can reliably capture the global folds of antibody fragments^26,27^.

**Figure 4.**
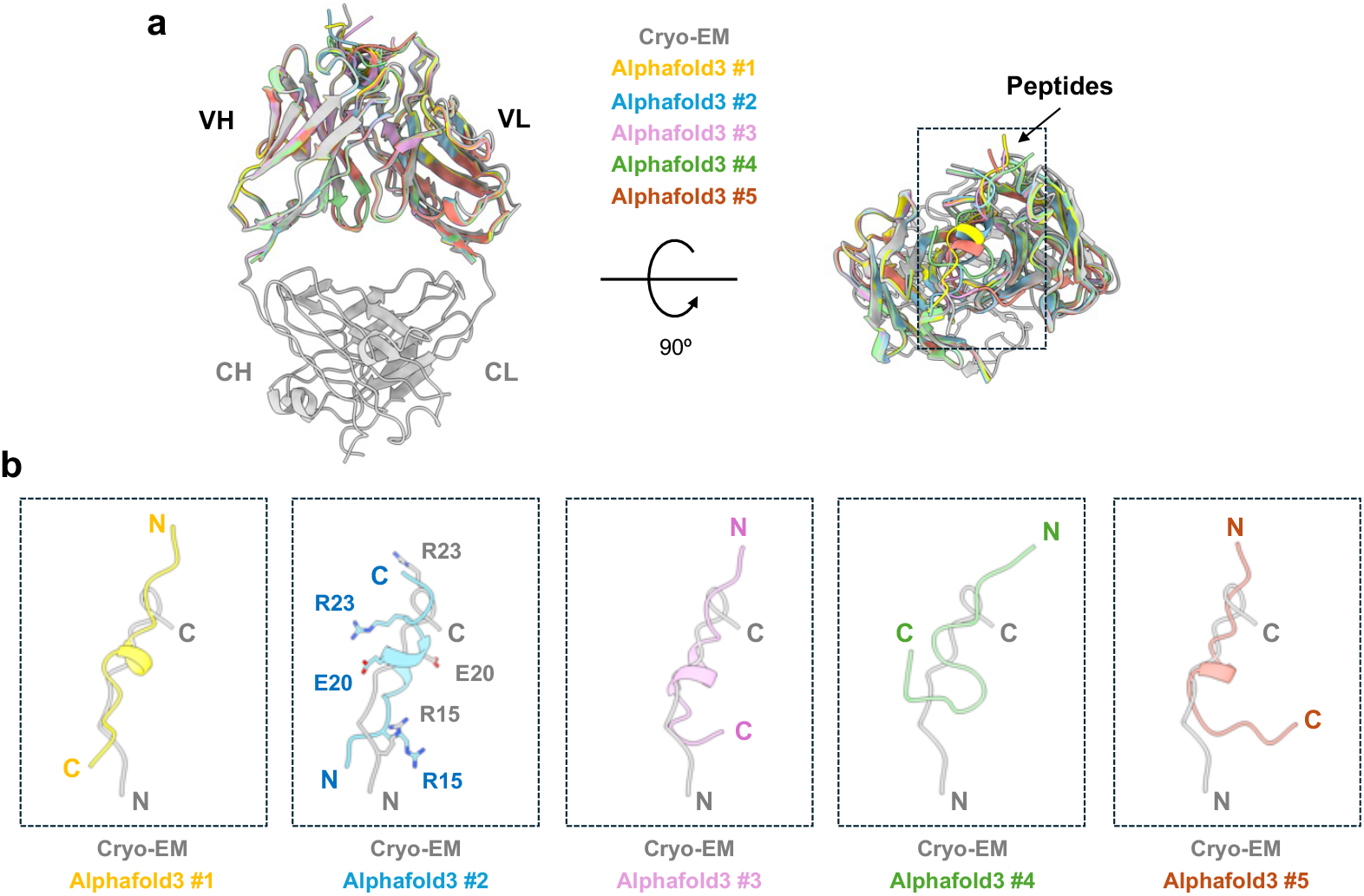
Cryo-EM versus AlphaFold models of 37E5 Fab–peptide complex. (a) Structural alignment of the cryo-EM structure of the 37E5 Fab (*gray*) with five AlphaFold3-predicted models, showing close agreement in the Fab framework regions. (b) Pairwise structural alignments between the cryo-EM structure and each of the five AlphaFold3 models.

Closer inspection of the binding interface revealed major discrepancies between the AlphaFold-predicted and experimentally determined structures. In the cryo-EM model, the MLC1-derived peptide was clearly resolved within the paratope, with well-defined side-chain densities that enabled mapping of salt bridges and hydrogen bonds (Fig. 2d, 3a, and 3b). In contrast, the AlphaFold prediction failed to position the peptide correctly within the binding pocket, and the modeled side chains were highly inaccurate and inconsistent with the observed interaction network in cryo-EM (Fig. 4b). Among the five predicted models, four misassigned the peptide orientation—swapping the amino- and carboxyl-terminal ends—while one model (AlphaFold3 #2) placed the peptide in the correct orientation but mislocalized several key side chains. As a result, critical epitope residues such as Arg15′, Glu20′, and Arg23′ were not oriented appropriately to engage the CDR loops. Although AlphaFold provides accurate Fab backbone predictions, it fails to reproduce the geometry of peptide binding or side-chain interactions at the paratope. These findings highlight the limitations of current computational approaches and reinforce the essential role of cryo-EM in defining antibody–antigen interfaces with mechanistic accuracy.

## DISCUSSION

Here, we introduce 37E5, a monoclonal antibody that specifically detects MLC1 across species—including human, mouse, and chicken—in diverse contexts, from heterologous expression in mammalian cells to native expression in mouse and human brain tissue. These properties establish 37E5 as a versatile reagent for biochemical assays, cellular imaging, and histological analysis of MLC1 in both model animals and humans. Thus, beyond its immediate application to biochemical studies, 37E5 provides a critical foundation for developing reliable biomarkers and detection tools for MLC1 in megalencephalic leukoencephalopathy with subcortical cysts (MLC).

In this study, using a conventional 300 kV cryo-EM system equipped with a Falcon 4 direct electron detector and a Selectris-X energy filter, we obtained ∼3 Å reconstructions of both apo and antigen-bound 37E5 Fab without the need for specialized instrumentation. The high local resolution enabled side-chain assignments and direct visualization of the epitope–paratope interface. These results demonstrate that careful optimization of sample preparation and data processing on standard cryo-EM platforms can push the lower size limits of cryo-EM, broadening its applicability to antibody fragments and other small protein complexes^7^.

The cryo-EM structures reveal how 37E5 achieves high specificity for MLC1. A network of polar contacts, involving Arg15′, Glu20′, Gly22′, and Arg23′ in the antigen peptide, together with complementary residues in the CDR loops, anchor the peptide within the binding cleft. Hydrophobic contacts, particularly between Met16′ and Leu19′ in MLC1 and aromatic residues in the Fab, further stabilize the complex. Notably, the conformational rearrangements of CDR-L1 and CDR-L2 upon antigen engagement demonstrate an induced-fit mechanism of recognition, in contrast to static lock-and-key models. This dynamic adaptability may represent a general feature of antibody recognition, enabling antibodies to fine-tune binding to linear epitopes presented in different structural contexts^28-31^.

By defining the structural and biochemical basis of 37E5-MLC1 recognition, our study provides a framework for rational antibody engineering. Detailed mapping of polar and hydrophobic interactions within the paratope allows identification of critical residues that determine binding specificity. Such information can be leveraged to engineer antibodies with altered affinity or species cross-reactivity, tailored to meet experimental, therapeutic, or diagnostic applications.

Although recent reports have suggested that AlphaFold models can predict protein–protein and protein–ligand interactions, including antibody–antigen recognition, with high accuracy^27,32,33^, the comparison of 37E5 Fab-peptide cryo-EM structure with AlphaFold3 predictions highlights both the promise and the current limitations of computational modeling for antibody–antigen complexes. While the Fab backbone was predicted with high accuracy, the peptide was misplaced within the binding pocket, and side-chain orientations failed to recapitulate the experimentally observed interaction networks. These discrepancies illustrate that current AI-based approaches, while powerful for predicting global protein folds, lack the accuracy to capture dynamic and context-dependent features of antibody– antigen recognition. Thus, our results reinforce the continued necessity of high-resolution experimental approaches, such as cryo-EM, to define antibody–antigen interfaces at atomic detail.

Collectively, our work establishes 37E5 as a valuable reagent for MLC1 research, defines the molecular determinants of MLC1 recognition, and demonstrates the feasibility of cryo-EM to resolve dynamic features of small Fab–antigen complexes. These findings advance both methodological and mechanistic understanding, with implications extending to antibody engineering, neurovascular biology, and rare leukodystrophy research. Future studies should leverage 37E5 to dissect the physiological role of MLC1 in gliovascular signaling and to evaluate potential therapeutic strategies for MLC disease. Looking forward, integration of cryo-EM data with AI-based predictions may offer a synergistic strategy, where experimental validation corrects and refines computational models, leading to improved predictive frameworks for antibody design.

## METHODS

### Immunization and hybridoma screening

Immunization was performed as previously described^16^. Female BALB/c mice (8 weeks old) were immunized intraperitoneally (i.p.) with 200 μL of a mixture containing 100 μg of KLH-conjugated peptide (N-YDRMPTLERGRQDA-C-KLH, AbClon) and 100 μL of Complete Freund’s Adjuvant (CFA, Sigma-Aldrich #F5881). For subsequent booster injections (three times), Incomplete Freund’s Adjuvant (IFA, Sigma-Aldrich #F5506) was used. Injections were given every 2 weeks. After the third i.p. injection, serum was collected and tested to assess immune responses against the peptide using ELISA. Pre- and post-immune sera were serially diluted (1:500∼1:10,000 (*v/v*)) and added to 96-well plates pre-coated with peptide (5–10 μg/mL BSA-conjugated peptide (N-YDRMPTLERGRQDA-C-BSA, AbClon) in PBS containing 2 mg/mL BSA). After 1 h incubation, plates were washed with PBS and incubated with goat anti-mouse IgG antibody conjugated to alkaline phosphatase (Sigma-Aldrich #A3562). After washing, bound antibodies were quantified using a *p*-Nitrophenyl Phosphate (PNPP) substrate kit (Thermo #37620), following the manufacturer’s instructions. Fifty μL of PNPP substrate/diethanolamine buffer mixture was added per well, and absorbance at 405 nm was measured. After confirming the immune response, mice received an additional booster immunization and were sacrificed for spleen isolation.

Hybridomas were generated by fusion of splenocytes with SP2/0 myeloma cells (ATCC #CRL-1581) as previously described^16^. Isolated splenocytes were resuspended in fusion medium (10% (*v/v*) NCTC medium (Sigma-Aldrich #N1140), 2 mM L-glutamine (Gibco #25030), 1 mM sodium pyruvate (Gibco #11360), 1x MEM NEAA (Gibco #11140), and 50 μg/mL gentamycin (Gibco #15750-078) in high-glucose DMEM (Gibco #11965092)) and mixed with SP2/0 cells at a 10:1 ratio. After removal of medium, 1 mL of 50% PEG 1500 solution (Roche #10 783 641 001) was carefully added dropwise. PEG was washed out with fusion medium, and cells were pelleted by centrifugation. FBS (Gibco #16000-044) was added to the pellet and incubated for 5 min at 37 °C. Cells were resuspended in complete medium (fusion medium supplemented with 10% FBS, 1x HAT supplement (Gibco #21060017), and 1x Hybridoma Fusion and Cloning Supplement (HFCS, Roche #11 363 735 001)) and seeded at 2.5 x 10^5^ cells/well in 96-well plates. Hybridoma culture supernatants were screened by PNPP-based ELISA to assess relative antibody production. Several rounds of selection were performed, and monoclonal hybridomas were isolated by limiting dilution and expanded for antibody purification. All animal procedures were approved by the Institutional Animal Care and Use Committee of the Korea Brain Research Institute (IACUC-23-00049 and IACUC-23-00048).

### Purification of 37E5 anti-MLC1 antibody

The 37E5 hybridoma clone was cultured in Hybridoma-SFM (Gibco #12045076) supplemented with 4% (*v/v*) ultra-low IgG FBS (Gibco #16250-078). Whole IgG was purified from culture supernatants by affinity chromatography using Protein G resin (GenScript #L00209). The resin was washed with 1x PBS (pH 7.4), and bound antibody was eluted with 0.1 M glycine (pH 2.2). Eluted fractions were immediately neutralized by the addition of 1.0 M Tris-HCl (pH 8.0) at a 1:1 volume ratio^25^.

### Western blotting and immunoprecipitation

HEK293T cells were maintained in DMEM supplemented with 10% FBS at 37 °C in a humidified incubator with 5% CO_2_. Cells were transfected with pcDNA3.1-Flag-human MLC1, pcDNA3.1-Flag-mouse Mlc1, or empty vector. For each transfection, 1 μg of DNA was mixed with 6 μg of polyethylenimine (PEI; PolyScience #24765-1) in Opti-MEM (Thermo Fisher #31985-062). After 48 h, cells were washed with 1x PBS (pH 7.4) and resuspended in lysis buffer (1% (*v/v*) Triton X-100 in PBS, pH 7.4), supplemented with protease inhibitor cocktail (Roche #05056489001). Lysates were incubated for 1 h at 4 °C and clarified by centrifugation. For immunoprecipitation, cleared supernatants were incubated with 37E5 antibody pre-bound to Protein G resin. After incubation for 16 h at 4 °C, unbound proteins were removed by washing with lysis buffer, and bound complexes were eluted with 2x lithium dodecyl sulfate (LDS) sample buffer (Thermo Fisher #B0007). Immunoblotting was performed using the iBlot and iBind systems (Thermo Fisher). The 37E5 antibody (1 μg/mL) was used as the primary antibody, and HRP-conjugated goat anti-mouse IgG (H+L) secondary antibody (0.2 μg/mL; Bio-Rad #170-6516) was used for detection^25^.

### Immunofluorescence staining of transfected cells

COS-7 cells were maintained in DMEM supplemented with 10% FBS at 37 °C in a humidified incubator with 5% CO_2_. For transient transfection, 1 μg of pCAG-Flag-hMLC1-GFP plasmid was used as above. After 48 h, cells were washed with PBS and fixed with 4% (*w/v*) paraformaldehyde (PFA) for 15 min at room temperature. Following removal of PFA, cells were washed with 1× phosphate-buffered saline (PBS) (pH 7.4), permeabilized with 0.2% (*v/v*) Triton X-100 in PBS, and washed again. Before antibody incubation, cells were incubated in blocking solution (3% (*w/v*) BSA and 0.1% (*v/v*) Tween-20 in PBS, pH 7.4). Primary antibodies, mouse anti-MLC1 (37E5), mouse anti-Flag (M2; Sigma #F1804), and chicken anti-GFP (Thermo Fisher #A10262) were diluted in blocking solution (0.4–1.0 μg/200 μL) and applied for 1 h at room temperature. After washing with wash solution (0.1% (*v/v*) Tween-20 in PBS), cells were incubated with secondary antibodies: donkey anti-mouse IgG (H+L)-Alexa Fluor 568 and goat anti-chicken IgY (H+L)-Alexa Fluor 488, each diluted in blocking solution (0.2 μg/200 μL). After removal of unbound antibodies, cells were mounted with ProLong Gold Antifade Mountant (Thermo Fisher #P36981). Images were acquired using a Nikon ECLIPSE Ti-E confocal microscope equipped with a Plan Apo 60×/NA 1.40 oil-immersion objective lens^18^.

### Human brain tissue

Postmortem human brain tissue was obtained from the hippocampus of an 80-year-old female donor who died of pneumonia, with a postmortem interval of 24 hours. Written informed consent for the use of cadavers was obtained from all donors or their authorized representatives, in strict adherence to established legal and ethical standards. All procedures were conducted in accordance with the Korean “Act on Anatomical Dissection and Preservation of Corpses” and the Declaration of Helsinki, with approval from the Institutional Review Board of the College of Medicine, The Catholic University of Korea (MC23EISI0014). The collected tissue was immediately washed with cold Dulbecco’s Phosphate-Buffered Saline (D-PBS, pH 7.1, Gibco, Frederick, Maryland, USA) to remove residual blood and then fixed in 4% (*w/v*) PFA in 0.1M phosphate buffer (PB, pH 7.4) for 24 h at 4°C. Subsequently, the tissue was cryoprotected in OCT medium and frozen for histology.

### Mouse brain tissue

All the animal experiments for immunohistochemistry were conducted in accordance with the animal care guidelines issued by the National Institutes of Health and by the Institutional Animal Use and Care Committee at the Catholic University of Korea (Approval number: CUMS-2024-0054-03). Mlc1 knockout (KO) and wild-type (WT) mice were generated by crossing Mlc1 STOP-tetO knock-in mice and tetO knock-in mice, respectively^23^. Genotyping and confirmation of MLC1 deletion were performed as previously described^34^. At 12 weeks of age, mice were anesthetized and perfused transcardially with cold 4% PFA in 0.1M PB, pH 7.4. The brains were removed and postfixed in 4% PFA overnight, then cryoprotected in 30% sucrose in 0.1 M PB.

### Immunofluorescence staining of brain tissues

Free-floating cryostat sections (25 μm) of brain tissue were blocked with blocking solution (0.2% gelatin, 1% bovine serum albumin, and 0.05% saponin in 0.01M PB) and incubated overnight at 4°C with primary antibodies: rabbit anti-MLC1 (1:100; Aviva Systems Biology, #ARP35249_P050), mouse anti-MLC1 (1 μg/mL; 37E5), and chicken anti-GFAP (1:500; Chemicon, #AB5541). Sections were subsequently incubated with secondary antibodies: Cy3-conjugated donkey anti-mouse (1:500; Jackson ImmunoResearch, #715-165-151), Cy3-conjugated donkey anti-rabbit (1:500; Jackson ImmunoResearch, #711-165-152), and Alexa Fluor 488–conjugated donkey anti-chicken (1:500; Jackson ImmunoResearch, #703-545-155). Cell nuclei were counterstained with DAPI (50 μg/mL; Roche, #10-236-276-001) for 15 min. Images were acquired using a confocal microscope (FV3000; Olympus).

### Fab preparation

Fab preparation was performed as previously described^16^. Briefly, purified 37E5 IgG (5 mg/mL) was digested with papain (50 μg/mL; Roche #10 108 014 001) in cutting buffer (100 mM NaCl, 5 mM EDTA, 10 mM MOPS, pH 7.0) supplemented with freshly prepared 20 mM L-cysteine for 1–2 h at 37 °C. The pH was readjusted to 7.0 after the addition of L-cysteine. The reaction was terminated by the addition of 30 mM iodoacetamide (freshly dissolved in H_2_O) and subsequently diluted ∼10-fold with low-salt HQ buffer (10 mM NaCl, 10 mM Tris-Cl, pH 8.0). The digested antibody was loaded onto a HiTrap Q HP anion-exchange chromatography column (Cytiva #17115301). The flow-through fraction (Fab) was collected and concentrated using Amicon Ultra centrifugal filter, 30 kDa MWCO (Millipore #UFC9030), whereas the bound fraction (Fc) was eluted with high-salt HQ buffer (400 mM NaCl, 10 mM Tris-Cl, pH 8.0). The Fab fraction was further purified using a Superdex 200 size-exclusion chromatography (SEC) column pre-equilibrated with 1x PBS (pH 7.4).

### Cryo-EM sample preparation and data collection

For the apo Fab structure, purified Fab (1–2 mg/mL, 3 μL) was applied to glow-discharged UltrAuFoil R1.2/1.3 300-mesh Au grids (SPI Supplies) or Quantifoil R1.2/1.3 300-mesh Cu grids (SPI Supplies) coated with a graphene oxide support film (Sigma-Aldrich). For the Fab–peptide complex, purified Fab (1–2 mg/mL) was incubated with peptide at a molar ratio of 1:1.2 prior to grid preparation. Grids were blotted for 5 s with blot force 5 using humidity-saturated Whatman No. 1 filter paper (Cytiva) and plunge-frozen in liquid ethane using a Vitrobot Mark IV (Thermo Fisher Scientific).

Cryo-EM data were collected on a Titan Krios G4 microscope (Thermo Fisher Scientific) operated at 300 kV, equipped with a Falcon 4 direct electron detector and a Selectris X energy filter (slit width 10 eV). Micrographs were recorded at a nominal magnification of 165,000x, corresponding to a calibrated pixel size of 0.7451 Å, with a total electron dose of 50 e^-^/Å^2^ and a defocus range of –0.6 to – 2.2 μm.

### Data processing and structure determination

Image processing was performed using cryoSPARC v4.4.0^35^. For all datasets, cryo-EM movies were motion-corrected with Patch Motion Correction, and contrast transfer function (CTF) parameters were estimated with Patch CTF Estimation. Micrographs were curated based on astigmatism and CTF fit resolution.

For the apo Fab dataset, particles were initially selected using Blob Picker, yielding 937,094 particles from 6,118 micrographs (UltrAuFoil) and 5,886,251 particles from 5,077 micrographs (graphene oxide–treated grids). These particles were extracted with a box size of 300 pixels and subjected to multiple rounds of 2D classification and Topaz^36^, resulting in 912,989 particles. Ab initio reconstruction was generated from this subset and classified into five distinct 3D classes, of which two classes comprising 414,528 particles were selected for further refinement. Non-uniform refinement in cryoSPARC^37^ yielded cryo-EM maps at 3.16 Å resolution.

For the Fab–peptide dataset, particle selection by Blob Picker yielded 1,913,958 particles from 3,662 micrographs (UltrAuFoil) and 6,146,011 particles from 4,902 micrographs (graphene oxide– treated grids). Extracted particles (300-pixel box size) were refined through 2D classification and Topaz, resulting in 568,838 particles. Ab initio reconstruction was performed and classified into six distinct 3D classes, from which two classes comprising 219,979 particles were selected for refinement. Non-uniform refinement yielded cryo-EM maps at 3.04 Å resolution.

### Atomic model building and refinement

For initial atomic models of the Fab variable regions and the antigen peptide, we used ModelAngelo (RELION 5)^24^ with the corresponding sequence information. For the Fab constant regions, we applied ModelAngelo without sequence input. The resulting models were fitted into the cryo-EM maps using ChimeraX^38^, manually rebuilt in COOT^39^, and refined with real space refinement in PHENIX^26,40^. Cryo-EM data collection, model refinement, and validation statistics are summarized in Table S1. Structural figures were prepared using ChimeraX.

## DATA AVAILABILITY

The final coordinates and cryo-EM maps supporting the findings of this study have been deposited in the Worldwide Protein Data Bank (www.wwpdb.org) under the following accession codes: PDB IDs (9VHV and 9X08) and EMDB IDs (EMD-65073 and EMD-66422).

## ACKNOWLEDGEMENTS

This work was supported by the KBRI basic research program through Korea Brain Research Institute funded by the Ministry of Science and ICT (25-BR-01-02 and 25-BR-05-07 to H.H.Lim) and by the ELA international (ELA no.2024-01814 to H.H.Lim). This research was also supported by Basic Medical Science Facilitation Program through the Catholic Medical Center of the Catholic University of Korea funded by the Catholic Education Foundation. We are grateful to Drs. R.Min (VU University medical center), J.Y.Moon (KBRI), and W.J.Oh (KBRI) for their initial evaluations for 37E5 antibody. We also thank all members of the Lim laboratory at KBRI for their timely help throughout the study.

## AUTHOR INFORMATION

**Neurovascular Unit Research Group, Korea Brain Research Institute (KBRI), Daegu, Republic of Korea**

Junmo Hwang & Hyun-Ho Lim

**Structural Biology Division, Baobab AiBIO Co**., **LTD**., **Incheon, Republic of Korea**

Kunwoong Park & Kyoung Tai No

**Department of Anatomy, College of Medicine, The Catholic University of Korea, Seoul, Republic of Korea**

Tae-Ryong Riew

**Department of Pharmacology, College of Medicine, The Catholic University of Korea, Seoul, Republic of Korea**

Jeong-Sun Choi & Kyung Ok Cho

**Department of Medical Science, College of Medicine, Graduate School of The Catholic University of Korea, Seoul, Republic of Korea**

Tae-Ryong Riew & Kyung Ok Cho

**Institute of Convergence Science and Technology, Yonsei University, Incheon, Republic of Korea**

Kyoung Tai No

**Department of Brain Sciences, DGIST, Daegu, Republic of Korea**

Hyun-Ho Lim

### Contributions

J.H. developed 37E5 antibody, conducted hybridoma culture and mAb purification, prepared Fab fragment, and performed biochemical studies. K.P. conducted cryo-EM data collection, structure determination, and structure analysis. J.S.C., T.R.R., and K.O.C. performed immunohistochemistry of mouse and human brains. K.T.N. supervised the cryo-EM study. H.H.L. conceptualized and supervised the project, processed and analyzed the data, and wrote the paper with contributions from all authors.

