## supplementary information for "Cryo-EM structures of an anti-MLC1 Fab in apo and peptide-bound states reveal the structural basis of antigen recognition"

### Extended Data Table 1 . Cryo-EM data collection and refinement statistics.

| Structure | Fab apo state (PDB #9VHV) | | Fab + peptide complex (PDB #9X08) | |
|---|---|---|---|---|
| **Data collection and processing** | | | | |
| Dataset (grid type) | *Fab_Apo (UltrAufoil)* | *Fab_Apo (Graphene-Oxide)* | *Fab + Peptide (UltrAufoil)* | *Fab + Peptide (Graphene-Oxide)* |
| Microscope | FEI Titan Krios | FEI Titan Krios | FEI Titan Krios | FEI Titan Krios |
| Detector | Falcon 4 | Falcon 4 | Falcon 4 | Falcon 4 |
| Magnification | x165,000 | x165,000 | x165,000 | x165,000 |
| Voltage (kV) | 300 | 300 | 300 | 300 |
| Electron Exposure ($e^-$/Å$^2$) | 50 | 50 | 50 | 50 |
| Exposure rate ($e^-$/pix/s) | 5.31 | 5.32 | 5.27 | 5.30 |
| Defocus Range (μm) | -0.8 to -2.2 | -0.6 to -2.0 | -0.8 to -2.2 | -0.6 to -2.0 |
| Pixel size (Å) | 0.7451 | 0.7451 | 0.7451 | 0.7451 |
| Number of movies | 6,118 | 5,077 | 3,662 | 4,902 |
| Symmetry imposed | *C*1 | | *C*1 | |
| Initial particle images (no.) | 6,823,345 | | 8,059,969 | |
| Final particle images (no.) | 391,736 | | 205,482 | |
| Map resolution (Å) | 3.16 | | 3.04 | |
| FSC threshold | 0.143 | | 0.143 | |
| **Refinement** | | | | |
| Initial model used(PDB) | Modelangelo | | Modelangelo | |
| Composition (#) | | | | |
| Atoms | 3362 | | 3463 | |
| Amino acids | 431 | | 443 | |
| Ligands | - | | - | |
| RMSD bonds (Å) (# >4σ) | 0.003 | | 0.004 | |
| RMSD angles (°) (# >4σ) | 0.602 | | 0.597 | |
| Mean B-factors (Å$^2$) | | | | |
| Amino acids | 45.42 | | 52.27 | |
| Ligand | - | | - | |
| Ramachandran (%) | | | | |
| Favored | 95.76 | | 97.47 | |
| Allowed | 4.24 | | 2.53 | |
| Outliers | 0 | | 0 | |
| Rotamer outliers (%) | 1.33 | | 0.52 | |
| Clash score | 5.43 | | 6.01 | |
| C-beta outliers (%) | NA | | NA | |
| CaBLAM outliers | 1.67 | | 2.11 | |
| CC (mask) | 0.85 | | 0.82 | |
| MolProbity score | 1.68 | | 1.44 | |

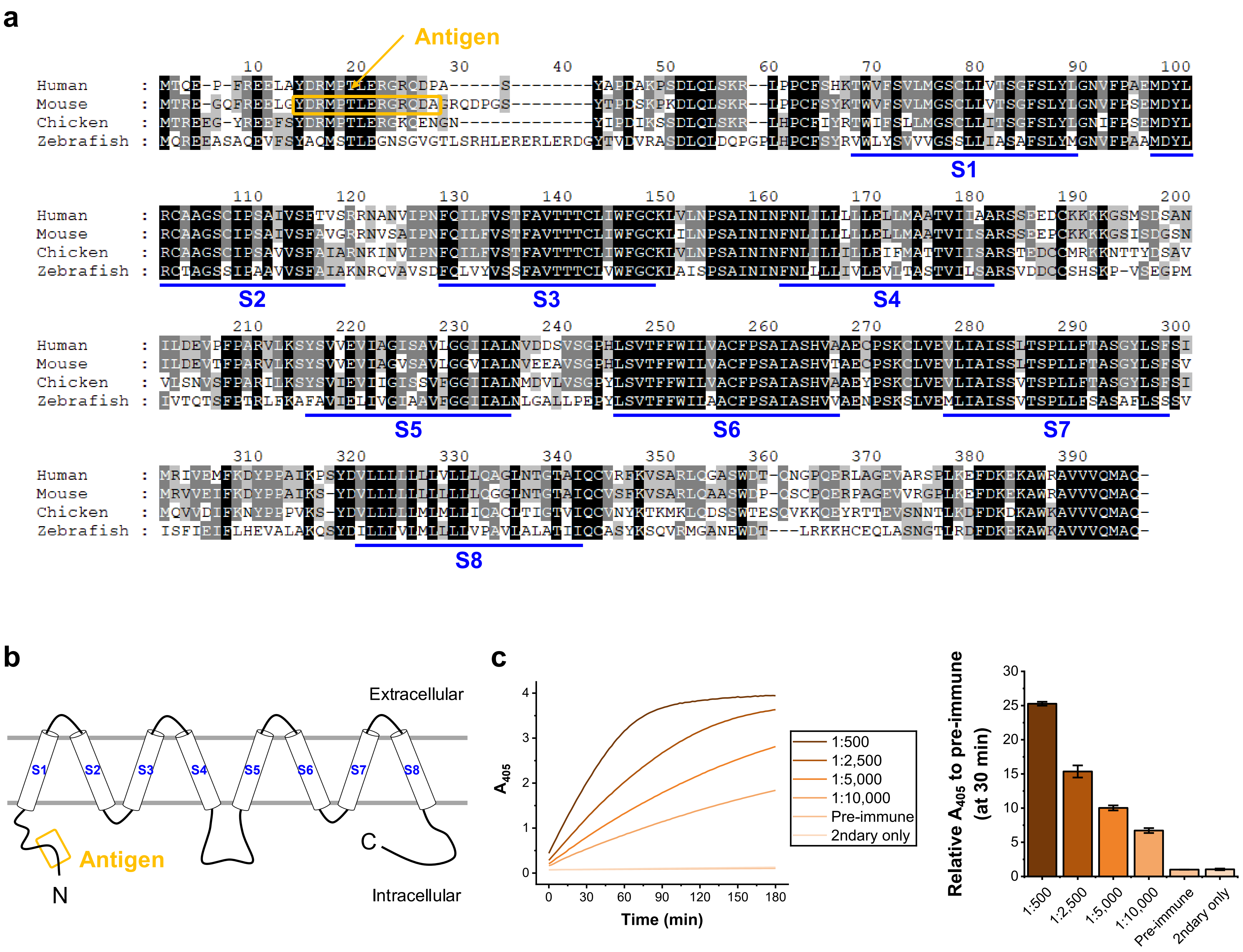


**Extended Data Figure 1. Sequence alignment and antigen region of MLC1 orthologs.**

(a) Multiple sequence alignment of MLC1 orthologs from human, mouse, chicken, and zebrafish (UniProt accession IDs: Q15049, Q8VHK5, A0A1D5PIT5, and A0A3P9CLQ4, respectively). Transmembrane domains (S1–S8) are indicated below the alignment. The antigen peptide sequence used for immunization is highlighted in yellow. (b) Schematic representation of the MLC1 transmembrane topology showing the position of the antigenic region. (c) ELISA results assessing the immune response against the antigen peptide following immunization.

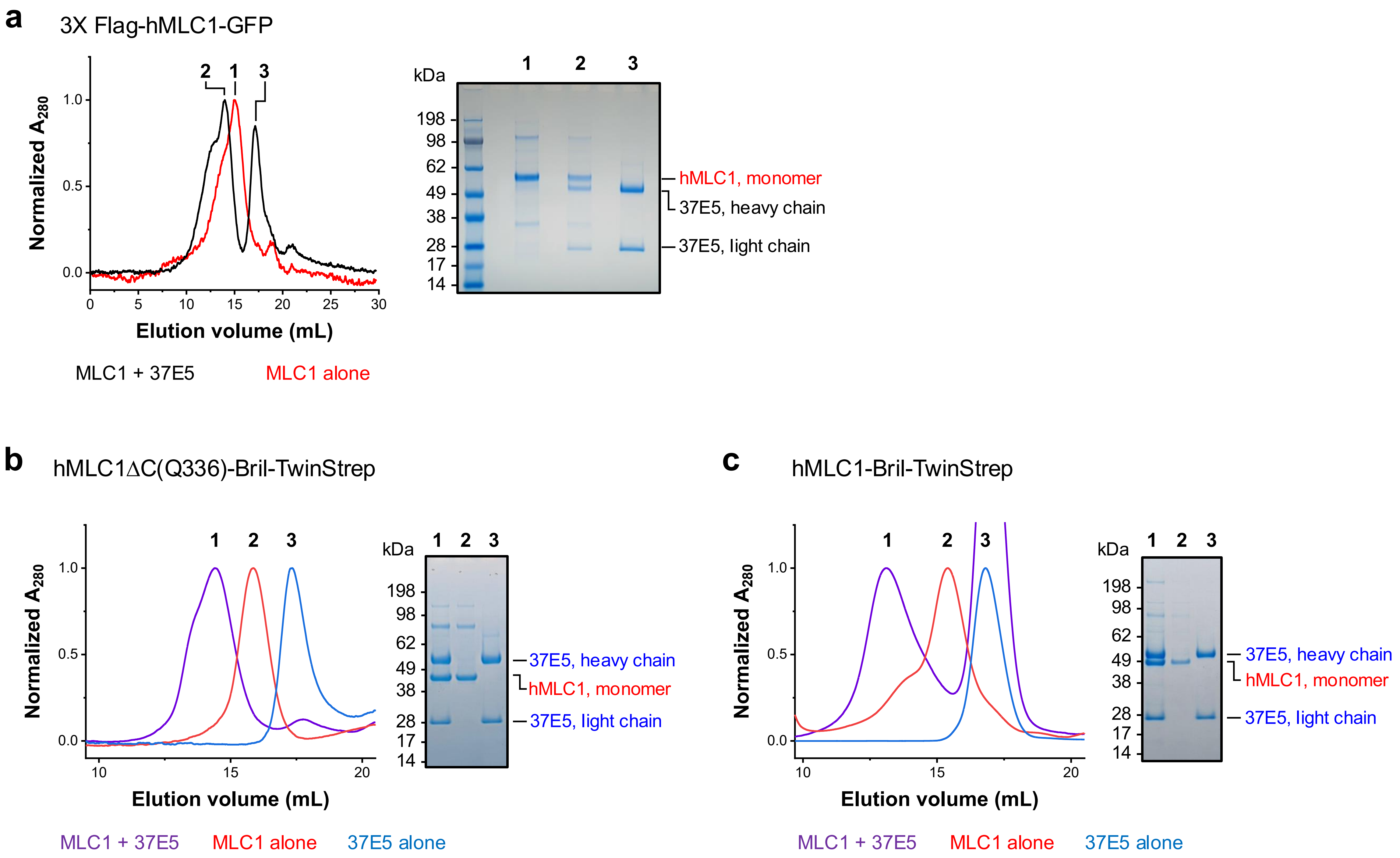


**Extended Data Figure 2. Binding of full-length 37E5 antibody to purified MLC1 proteins with various affinity tags.**

Size-exclusion chromatography (SEC) profiles and corresponding SDS–PAGE analyses showing the interaction between full-length 37E5 antibody and purified MLC1 proteins carrying different affinity tags: (a) 3×Flag–hMLC1–GFP, (b) hMLC1ΔC(Q336)–Bril–TwinStrep, and (c) hMLC1–Bril–TwinStrep. Co-elution of MLC1 with 37E5 antibody indicates stable complex formation in solution.

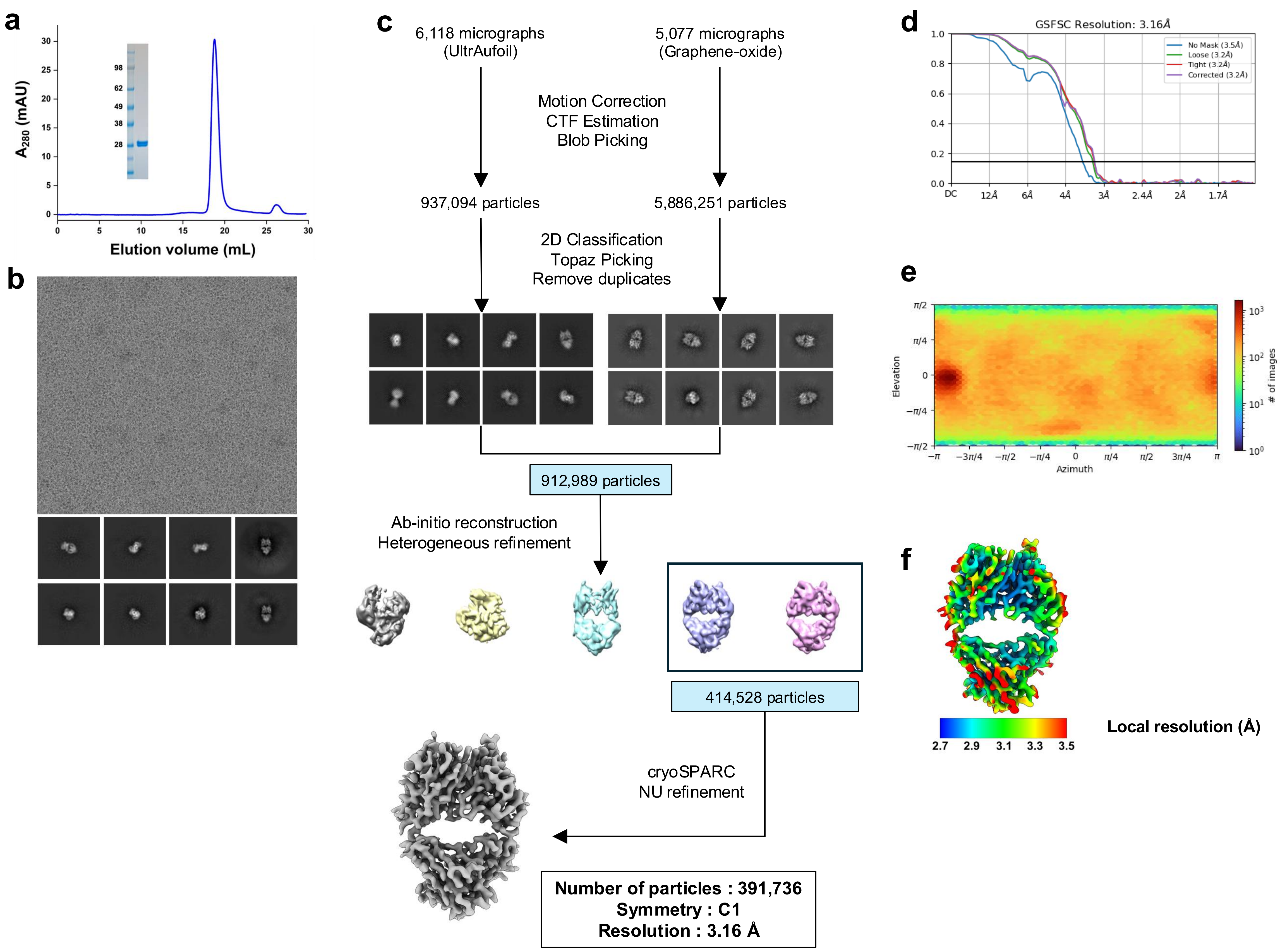


**Extended Data Figure 3 . Cryo-EM workflow for structural determination of the 37E5 Fab in the apo state.**

(a) Size-exclusion chromatography (SEC) profile of purified 37E5 Fab, showing a single monodisperse peak corresponding to the Fab fragment. (b) Representative cryo-EM micrograph and selected 2D class averages of Fab particles. (c) Image processing workflow used for structure determination, including particle picking, 2D classification, ab initio reconstruction, heterogeneous refinement, and non-uniform refinement in cryoSPARC. (d) Gold-standard Fourier shell correlation (GSFSC) curve used for global resolution estimation. (e) Angular distribution plot of particle orientations. (f) Local resolution map of the final cryo-EM reconstruction.

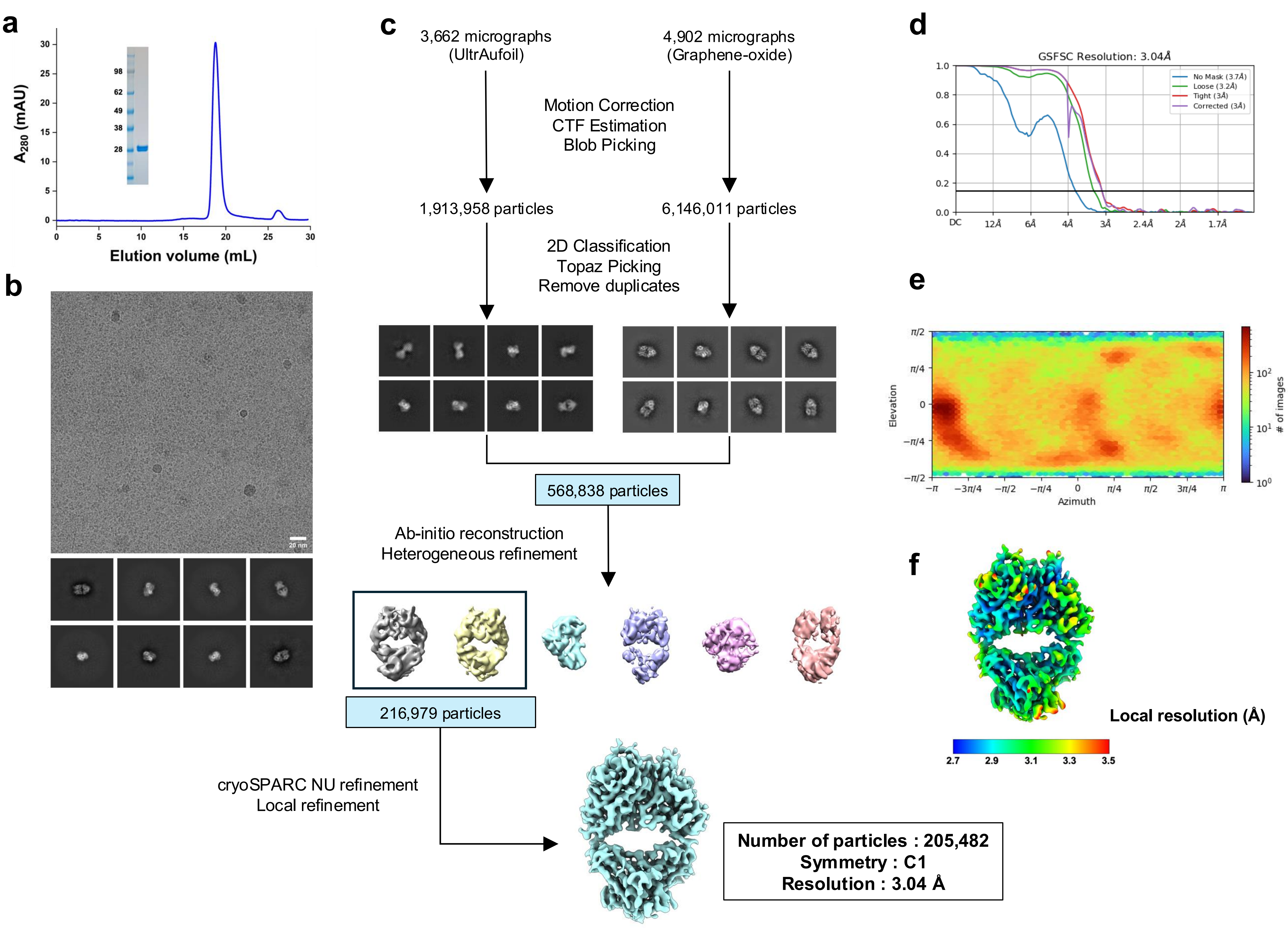


**Extended Data Figure 4 . Cryo-EM workflow for structural determination of the 37E5 Fab in the antigen-bound state.**
(a) Size-exclusion chromatography (SEC) profile of purified 37E5 Fab, showing a single monodisperse peak corresponding to the Fab fragment. (b) Representative cryo-EM micrograph and selected 2D class averages of Fab–peptide particles. (c) Image processing workflow used for structure determination, including particle picking, 2D classification, ab initio reconstruction, heterogeneous refinement, and non-uniform refinement in cryoSPARC. (d) GSFSC curve used for global resolution estimation. (e) Angular distribution plot of particle orientations. (f) Local resolution map of the final cryo-EM reconstruction.

### a. Heavy chain

FR 1 | CDR-H1 | FR 2 | CDR-H2

1 QIQLVQSGPDLKKPGETVKISCKASGYTFTNYGMSWVKQAPGKVLKWMGWINTYTGEPTY 60

FR 3 | CDR-H3 | FR 4

61 ADDFKGRFAFSLETSANTAYLQINNLKNEDTATYFCAREGATSGFPYWGQGTLVTVSAAK 120

121 TTAPSVYPLAPVCGDTTGSSVTLGCLVKGYFPEPVTLTWNSGSLSTGVYTFPAVLRDKHR 180

181 DLTASVTINSDNFGGTHELECNVACTSCDDKTERKINPNG 220

### b. Light chain

FR 1 | CDR-L1 | FR 2 | CDR-L2

1 DIVMSQSPSSLAVSAGEKVTMSCRSSQSLLNSRTRKNYLAWYQQKPGQSPKLLIYWASTR 60

FR 3 | CDR-L3 | FR 4

61 ESGVPDRFTGSGSGTDFTLTISSVQAEDLAVYYCKQSYDLPYTFGGGTKLEIKRADAAPT 120

121 VSIFPPSSEQLTSGGASVVCFLNNFYPKDINVKWKIDGSECKTGVEETWTEFNSKDSNYS 180

181 MCSTLTLTREEWKNWNSATCKATQNTETEPIIKSFIRDL 219

**Extended Data Figure 5. Amino acid sequences of the 37E5 Fab fragment.**

Amino acid sequences of the 37E5 Fab heavy (a) and light (b) chains. Residues in the variable regions are shown in blue, and those in the constant regions are shown in black. Complementarity-determining regions (CDRs) are highlighted with red boxes, and framework regions (FRs) are indicated in green.

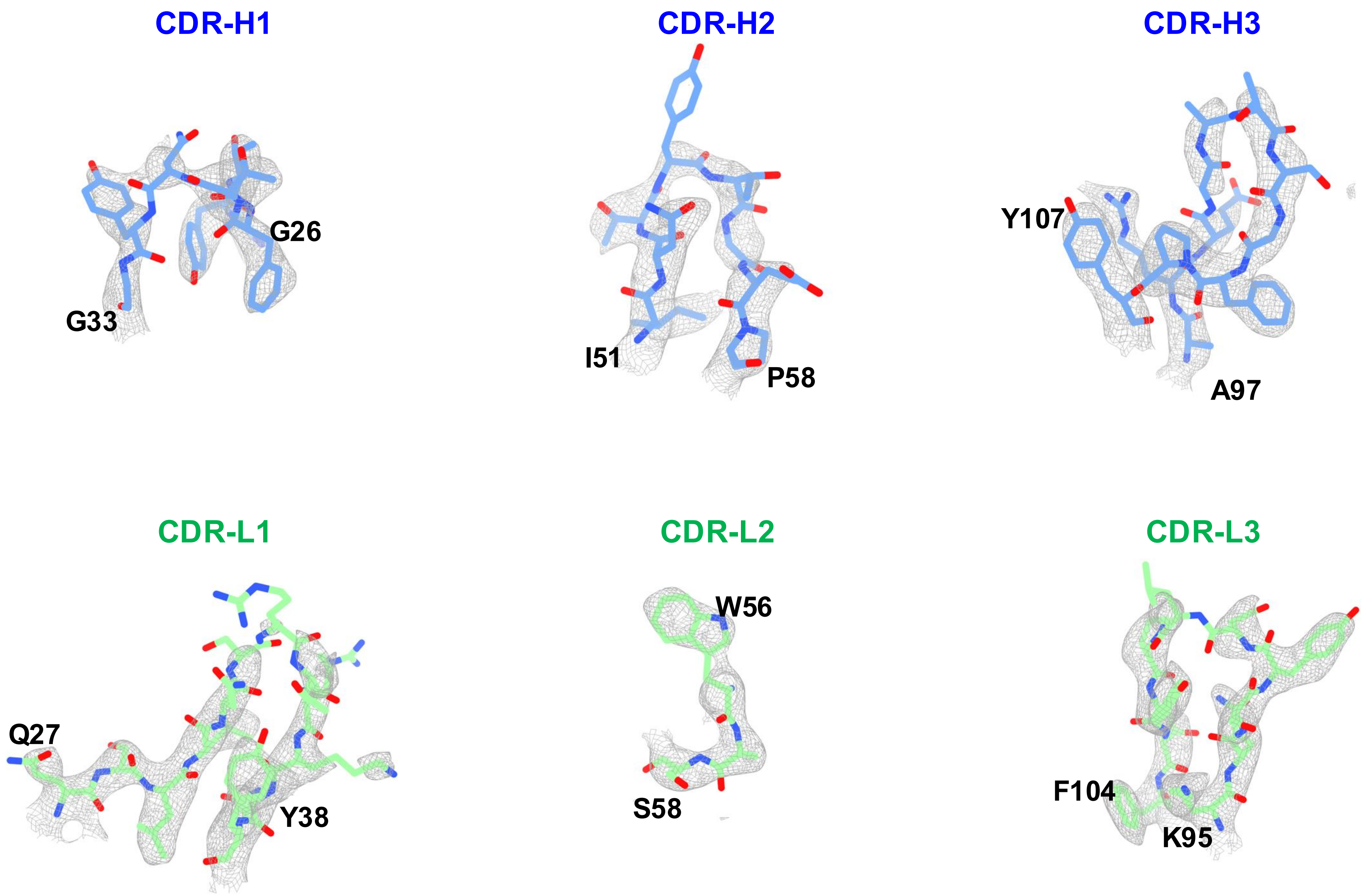


**Extended Data Figure 6 . Local density and atomic model of the apo Fab.**
Local density views of the CDR regions in the heavy and light chains, showing continuous and well-defined electron density that enables unambiguous side-chain visualization and accurate model fitting.

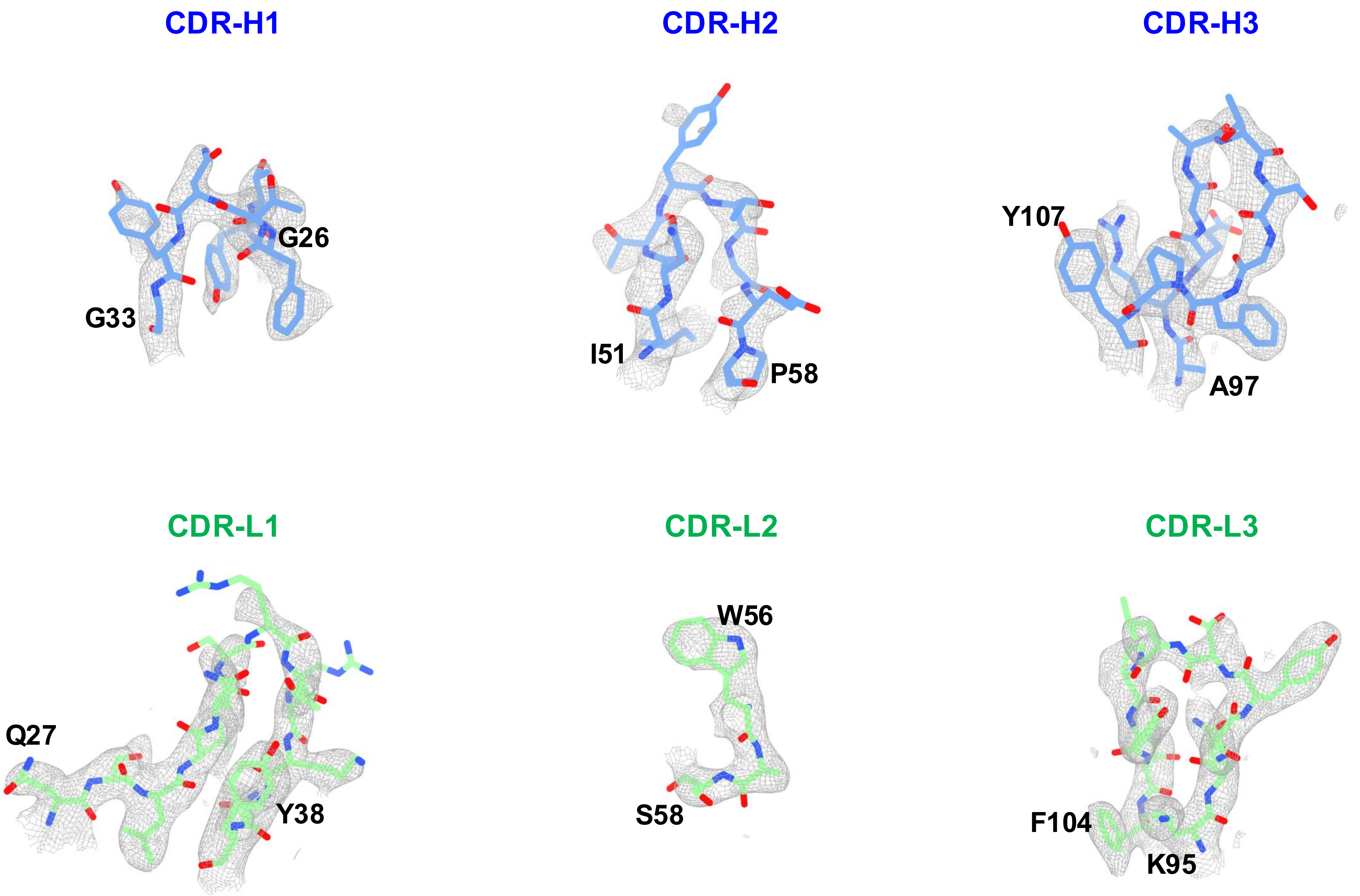


**Extended Data Figure 7 . Local density and atomic model of the Fab-peptide complex.** Local density views of the CDR regions in the heavy and light chains, showing continuous and well-defined electron density that enables unambiguous side-chain visualization and accurate model fitting.
